# A bovine model of rhizomelic chondrodysplasia punctata caused by a deep intronic splicing mutation in the *GNPAT* gene

**DOI:** 10.1101/2024.06.13.598642

**Authors:** Arnaud Boulling, Julien Corbeau, Cécile Grohs, Anne Barbat, Jérémy Mortier, Sébastien Taussat, Vincent Plassard, Hélène Leclerc, Sébastien Fritz, Cyril Leymarie, Lorraine Bourgeois-Brunel, Alain Ducos, Raphaël Guatteo, Didier Boichard, Mekki Boussaha, Aurélien Capitan

**Affiliations:** Université Paris Saclay, INRAE, AgroParisTech, GABI, 78350, Jouy-en-Josas, France; Oniris, INRAE, BIOEPAR, 44300, Nantes, France; Service d’imagerie médicale, DEPEC, Ecole Nationale Vétérinaire d’Alfort, 94700 Maisons-Alfort, France; ELIANCE, 75012 Paris, France; OS Aubrac, 12000 Rodez, France; GenPhySE, Université de Toulouse, INRAE, ENVT, 31326 Castanet Tolosan, France

## Abstract

**Background:** Genetic defects that occur naturally in livestock species provide valuable models for investigating the molecular mechanisms underlying rare human diseases. Livestock breeds are subject to the regular emergence of recessive genetic defects, due to their low genetic variability, while their large population sizes provide easy access to case and control individuals, as well as massive amounts of pedigree, genomic and phenotypic information recorded for selection purposes. In this study, we investigated a lethal form of recessive chondrodysplasia observed in 21 stillborn calves of the Aubrac breed of beef cattle.

**Results:** Detailed clinical examinations revealed proximal limb shortening, epiphyseal calcific deposits and other clinical signs consistent with human rhizomelic chondrodysplasia punctata, a rare peroxisomal disorder caused by recessive mutations in one of five genes (*AGPS, FAR1*, *GNPAT*, *PEX5* and *PEX7*). Using homozygosity mapping, whole genome sequencing of two affected individuals, and filtering for variants found in 1,867 control genomes, we reduced the list of candidate variants to a single deep intronic substitution in *GNPAT* (g.4,039,268G>A on Chromosome 28 of the ARS-UCD1.2 bovine genome assembly). For verification, we performed large-scale genotyping of this variant using a custom SNP array and found a perfect genotype-phenotype correlation in 21 cases and 26 of their parents, and a complete absence of homozygotes in 1,195 Aubrac controls. The g.4,039,268A allele segregated at a frequency of 2.6% in this population and was absent in 375,535 additional individuals from 17 breeds. Then, using *in vivo* and *in vitro* analyses, we demonstrated that the derived allele activates cryptic splice sites within intron 11 resulting in abnormal transcripts. Finally, by mining the wealth of records available in the French bovine database, we demonstrated that this deep intronic substitution was responsible not only for stillbirth but also for juvenile mortality in homozygotes and had a moderate but significant negative effect on muscle development in heterozygotes.

**Conclusions:** We report the first spontaneous large animal model of rhizomelic chondrodysplasia punctata and provide both a diagnostic test to counter-select this defect in cattle and interesting insights into the molecular consequences of complete or partial GNPAT insufficiency in mammals.

## Background

Over the last decade, the advent of high-throughput genotyping and next-generation sequencing have dramatically advanced clinical research, leading to the identification of thousands of disease-causing variants in humans and non-model species [1]. However, most of genetic studies are biased by a tendency to focus on the exome because of the challenges of annotating non-coding regions of the genome and, to a lesser extent, to the advantages of whole-exome sequencing over whole-genome sequencing [2]. While increasing evidence points to the role of non-coding variations in the onset of diseases [3], their study in humans suffers from several limitations and the pathophysiology of many disorders remains unresolved. The main limiting factors include the small number of patients affected by rare genetic defects and the relatively high genetic variability of our species, which makes it difficult to filter variants in the absence of functional annotation. In addition, the difficulty in obtaining a variety of tissues from living or deceased patients due to health risks, ethical or religious concerns often hinders in-depth clinical investigation and functional validation. In this context, naturally occurring genetic defects in livestock species represent valuable models to study the molecular mechanisms underlying rare human diseases. Indeed, farm animals are divided into numerous inbred populations or breeds that are prone to the regular emergence of recessive genetic defects [4]. In addition, their large population sizes provide easy access to case and control individuals, as well as massive amounts of pedigree, genomic and phenotypic information recorded for selection purposes [5,6].

From 2002 to 2020, 21 stillborn Aubrac calves with severe skeletal dysplasia were reported to the French National Observatory for Bovine Abnormalities (ONAB) for initial suspicion of Bulldog Calf Syndrome (BDS), a congenital form of bovine chondrodysplasia previously described to be associated with mutations in the aggrecan (*ACAN*), and collagen type II alpha 1 chain (COL2A1) genes [5,7–13]. Pathological examination and pedigree analysis revealed that this new genetic defect was actually similar to human rhizomelic chondrodysplasia punctata (RCDP) [14]. RCDP is a recessive peroxisomal disease caused by mutations in five genes: *AGPS*, *FAR1*, *GNPAT, PEX5* and *PEX7* encoding the alkylglycerone phosphate synthase, the fatty alcohol reductase 1, the glyceronephosphate O-acyltransferase, and the peroxisomal biogenesis factor 5 and 7, respectively [15–20].

In this article we describe, how we were able to reduce the list of candidate causative variants to a single deep intronic substitution in *GNPAT*, and to characterize its molecular and clinical consequences in the homozygous and heterozygous states, by exploiting the wealth of resources available in cattle. In other words, we report the first large animal model of RCDP and an original example of a deep intronic mutation responsible for a genetic defect in livestock.

## Methods

### Animals

Twenty-one stillborn calves (8 males and 13 females) affected by a severe form of skeletal dysplasia were observed in 21 purebred Aubrac herds over an 18-year period. Seventeen of the breeders kept genealogical records, which were extracted from the French national pedigree database. Veterinarians and artificial insemination (AI) technicians performed a gross clinical description in the field and collected ear biopsies and photographs for all affected calves. Due to the rapid removal of carcasses by rendering companies, only three affected calves could be recovered for full necropsy and pathological examination. The body of an unaffected calf that died of natural causes at four days of age was also collected to serve as a control. At the time of the study, biological samples for DNA extraction were also available for 15 dams and 11 sires of the cases and for one of their common ancestors, the AI bull “E.”. In addition, after identification of the *GNPAT* candidate causal variant, blood from three heterozygous mutant and three wild-type cows was collected in PAXgene Blood RNA Tubes (Qiagen) to perform RNA extractions and RT-PCR analysis. Finally, whole genome sequences, SNP array genotypes, and phenotypes from thousands to hundreds of thousands of animals from numerous breeds collected in the framework of other projects were also used in this study. Details of this additional material and on the analyses performed on it are given below.

### Pedigree analysis

Genealogical information was extracted from the French national pedigree database for 17 of the affected calves and 110,247 control calves born between 2019 and 2021 with both parents recorded. A search for common ancestry between the parents of the affected calves was performed using the anc_comm option of the pedig package [21]. In parallel, the genetic contribution to the case and control populations was estimated for each ancestor using the prob_orig option of the same package. Then, for individuals with a genetic contribution greater than or equal to 1% in each population, the ratio “contribution to the case population/contribution to the control population” was calculated.

### Necropsy and pathological examination

The frozen bodies of one control and three affected calves were subjected to digital radiography (XDR1, Canon Medical Systems), computed tomography (CT) scanning (80-slice CT scanner, Aquilion lighting, Canon Medical Systems) and necropsy at the National Veterinary School of Alfort. In addition to the classic post-mortem examination, special attention was given to the deformities of the skull and limb bones. The heads were sawed through the midline, and the left limbs were harvested, boiled, cleaned of residual soft tissue, and bleached with 5% hydrogen peroxide, prior to partial skeletal reconstruction.

### DNA extraction

Genomic DNA was extracted from blood using the Wizard Genomic DNA Purification Kit (Promega) and from ear biopsies or semen using the Gentra Puregene Cell and Tissue Kit (Qiagen). DNA purity and concentration were evaluated using a NanoDrop spectrophotometer (ThermoFisher Scientific).

### Homozygosity mapping

The twenty-one Aubrac cases, 26 of their parents and 1548 control animals from the same breed were genotyped with various Illumina SNP arrays over time (Bovine SNP50, EuroG10K and EuroGMD). Genotypes were phased and imputed to the Bovine SNP50 using FImpute3 [22] in the framework of the French genomic evaluation, as described in Mesbah-Uddin *et al.* [23]. The position of the markers was based on the ARS-UCD1.2 bovine genome assembly. We then considered sliding haplotypes of 20 markers (∼1 Mb) and we computed Fisher’s exact tests on 2×2 contingency tables consisting of the number of homozygous and “non-homozygous” animals in the case and control groups. A Bonferroni correction was applied to account for multiple testing (n=78861 tests with at least one homozygous carrier in the case group) and therefore the 0.05 significance threshold was set at −log P=6.20.

### Analysis of Whole Genome Sequences

The genomes of two RCDP-affected calves were sequenced at 19.2x and 19.4x coverage on an Illumina NovaSeq6000 platform in 150 paired-end mode, after library preparation using the NEXTflex PCR-Free DNA Sequencing Kit (Perkin Elmer Applied Genomics). Reads were aligned to the ARS-UCD1.2 bovine genome assembly [24] with the Burrows–Wheeler aligner (BWA-v0.6.1-r104; [25]) prior to the identification of SNPs and small InDels using the GATK-HaplotypeCaller software [26] as previously described in Daetwyler *et al.* and Boussaha *et al.* [26, 27]. Putative structural variations (SVs) were detected using the Pindel [28], Delly [29], and Lumpy [30] software and recorded if they were scored by at least two tools in the same individual. These variants were compared with those found in 1,867 control genomes in a previous study using the same procedure [6]. The control genomes included 39 Aubrac individuals, all of whom were non-carriers of the 35-marker haplotype common to affected calves based on phased and imputed Illumina BovineSNP50 array genotypes, and representatives of more than 70 cattle breeds or populations (Additional file 1: Table S1). Only SNPs, InDels and SVs located within the mapping interval (Chr28:3,555,723-5,143,700 bp), observed in the homozygous state in both cases and absent in all controls were considered as candidate variants. The only remaining variant after filtering, in this case variant g.4,039,268G>A on Chr28 were annotated using Variant Effect Predictor (Ensembl release 110; https://www.ensembl.org/info/docs/tools/vep/index.html) [31].

### Validation of the variant g.4,039,268G>A by Sanger sequencing and large-scale genotyping

As a first verification, we genotyped the variant g.4,039,268G>A using PCR and Sanger sequencing in 6 Aubrac cattle (2 affected calves, 2 heterozygous parents of cases, and two non-carriers based on haplotype information). A segment of 640 bp was amplified in a Mastercycler pro thermocycler (Eppendorf) using primers 5’-TCCCTTCCTTCAAGGCTACA-3’ and 5’-GTTAGGAGCCAGAGCAGCAC-3’ and the Go-Taq Flexi DNA Polymerase (Promega), according to the manufacturer’s instructions. Amplicons were purified and bidirectionally sequenced by Eurofins MWG (Hilden, Germany) using conventional Sanger sequencing. Electropherograms were analyzed using NovoSNP software for variant detection [32].

In addition, to genotype this variant on a large scale, we added a probe to the Illumina EuroGMD SNP array using the following design: TTTGTTCAGTAGGAAGTGAGGGCAGCCATTTTGAGCATAACATGATTCTCAGTGT TTTTC[A/G]NNCTTGCCGCATGCACTTTTGTTTAAATGTGAGGAGAGTATGGCTGTATACAAAGTGAAA. The EuroGMD SNP array is routinely used for genomic evaluation in France and genotypes of 21 affected calves and 376,730 controls from 19 French breeds (including 1,195 Aubrac cattle) were available at the time of writing.

### Minigenes construction

A 1149 bp fragment containing exon 11, intron 11 and exon 12 of the *GNPAT* gene was amplified from the genomic DNA of a homozygous carrier of the Chr28 g.4,039,268A mutant allele. BamHI and XhoI restriction sites were incorporated into primers 5’-TACCGAGCTCGGATCCTCCAGAGGATGTCTACAGTTGC-3’ and 5’-GCCCTCTAGACTCGAGTTGCAAAGATTTACACACCTGA-3’ designed for this purpose. PCR was performed in a 25 μl reaction mixture containing 12.5µL 2X KAPA HiFi HotStart ReadyMix (Roche), 50 ng genomic DNA, and 0.3 μM each primer. The PCR program comprised an initial denaturation at 95 °C for 3 min followed by 30 cycles of denaturation at 98 °C for 20 s, annealing at 65 °C for 15 s, extension at 72 °C for 1 min, and a final extension at 72 °C for 1 min 20 s. The PCR products were cloned into the pcDNA3.1(+) vector (Invitrogen) that was linearized by restriction enzymes BamHI and XhoI, using the T4 DNA Ligase (New England Biolabs) in accordance with the manufacturer’s instructions. The resulting minigene construct carrying the alternative g.4,039,268A allele was termed pcDNA3.1-GNPAT_A. The g.4,039,268G reference allele was then introduced into the pcDNA3.1-GNPAT_A minigene construct by site directed mutagenesis to obtain the pcDNA3.1-GNPAT_G minigene construct. This was achieved by means of the QuikChange II XL Site-Directed Mutagenesis Kit (Agilent) using primers 5’-CTCAGTGTTTTTCGGACTTGCCGCATGC-3’ and 5’-GCATGCGGCAAGTCCGAAAAACACTGAG-3’ in accordance with the manufacturer’s instructions. Sequences of both minigenes were verified by Sanger sequencing using T7 and BGH universal primers in addition to primer 5’-CAAGTGGGTCTGGGGTCTG-3”.

### Cell culture and transfection

Human embryonic kidney (HEK) 293T cells were maintained in DMEM supplemented with 10% fetal calf serum (Gibco). Cells were seeded with 300 000 cells/well in 6-well plates and transfected 24 hours later with 1 µg of each minigene construct mixed with 3 µL of Lipofectamine 2000 Reagent (Invitrogen) per well, according to the manufacturer’s instructions. Four hours after transfection, media were replaced by DMEM supplemented with 10% fetal calf serum and maintained in an incubator at 37°C and 5% CO2. Forty-height hours after transfection, the cells were washed with phosphate-buffered saline (PBS) and lysed with RLT buffer (Qiagen).

### RNA extraction from cell culture and RT-PCR analysis

Total RNA was extracted from lysed transfected cells by means of the RNeasy Mini Kit according to the manufacturer’s instructions (Qiagen). The RT step was performed using the SuperScript® III First-Strand Synthesis System for RT-PCR (Invitrogen) with 400 ng RNA, OligodT20 5µM, 500 µM each dNTP, MgCl2 5 mM, DTT 0.01M, 40 U RNaseOUT and 200 U Superscript 3 following the manufacturer’s instruction. RT products were treated with 2U RNase H during 20 min at 37°C. The PCR step was achieved using primers forward T7 (5’-TAATACGACTCACTATAGGG-3’) and reverse BGH (5’-TAGAAGGCACAGTCGAGG-3’) located within the 5’- and 3’-untranslated regions of pcDNA3.1-GNPAT minigene constructs, respectively. The reaction was performed in a 50-μL mixture containing 1.25 U GoTaq DNA polymerase (Promega), 200 μM dNTPs, 2 μL cDNA and 0.5 μM of each primer. The PCR program had an initial denaturation at 95°C for 2 min, followed by 30 cycles of denaturation at 95°C for 30 s, annealing at 55°C for 30 s, extension at 72°C for 1 min 30 s, and a final extension step at 72°C for 5 min.

### RNA extraction from blood and RT-PCR analysis

RNA was extracted from blood samples collected in PAXgene Blood RNA Tubes using the PAXgene Blood RNA Kit (Qiagen). Tubes were stored at −20°C for one month and thawed at room temperature for 2 hours before RNA extraction. Next, they were centrifuged 10 min at 4000 g and the supernatant was gently discarded by pouring off the tube. RNA was extracted from cell pellets by following the manufacturer’s guidelines, and their purity and integrity were assessed with the Bioanalyzer 2100 (Agilent). A 1 µg commercial sample of bovine muscle RNA (Gentaur) was used as a positive control. The reverse transcription (RT) step was performed using the SuperScript III First-Strand Synthesis System for RT-PCR (Invitrogen) with 60 ng RNA, OligodT20 5µM, 500 µM each dNTP, MgCl_2_ 5 mM, DTT 0.01M, 40U RNaseOUT and 200 U Superscript 3 following the manufacturer’s instruction. RT products were treated with 2U RNase H for 20 min at 37°C. The PCR step was achieved using primers 5’-GCTTTCGCTTCCTATGCAGT-3’ and 5’-TGTCCCTCGTCATCACTTGT-3’ located within the *GNPAT* exon 11 and 12, respectively. Each cDNA sample was amplified in quadruplicate in a 25-μL mixture containing 0.75 U GoTaq DNA polymerase (Promega), 200 μM dNTPs, 1 μL cDNA and 0.5 μM of each primer. The PCR program had an initial denaturation at 95°C for 2 min, followed by 45 cycles of denaturation at 95°C for 30 s, annealing at 53°C for 30 s, extension at 72°C for 1 min, and a final extension step at 72°C for 10 min. The four PCR replicates obtained from each sample were pooled and concentrated with the MinElute PCR Purification Kit (Qiagen) before gel electrophoresis.

### Prediction of exonic splicing enhancers (ESE) and protein structure analysis

ESE motif prediction was performed using ESEfinder 3.0 software in the context of the A and G alleles to identify the creation or disruption of putative splicing regulatory elements (http://krainer01.cshl.edu/cgi-bin/tools/ESE3/esefinder.cgi?process=home; [33,34]). The coding sequence of normal and abnormal GNPAT transcripts was translated into amino acid sequences using ExpASy (https://web.expasy.org/translate/; [35]). Information about protein domains was obtained from UniProt (https://www.uniprot.org/uniprotkb/A4IF87/entry, accessed 08.05.2020) and from Ofman *et al.* [18].

### Effects of allele Chr28 g.4,039,268A on juvenile mortality rates

The phenotypic effects of four mating types on juvenile mortality rates were evaluated using records from the French national bovine database and the following fixed-effect model:

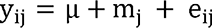

where yij represents the phenotype of interest, μ is the overall phenotypic mean, mj is the fixed effect of the mating status, and eij is the random residual error. The analysis was performed using the GLM procedure of SAS software (version 9.4; SAS Institute Inc., Cary, NC). The mating types considered were named 1 x 1, 1 x 0, 0 x 1, and 0 x 0, where the first position corresponds to the genotype of the sire and the second position corresponds to the genotype of the maternal grandsire in terms of allele dosage for the g.4,039,268A allele. Juvenile mortality was examined during four periods commonly considered in the literature (0-2, 3-14, 15-55 and 56-365 days after birth; e.g. [36,37]) as well as for a combination of the first two periods. For each time window, the mortality rates were calculated by dividing the number of calves that died of natural causes during the period by the number of calves present at the beginning of the period. We also calculated the expected effect on juvenile mortality rates, assuming a full penetrance of lethality under homozygosity, using the formula 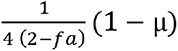 adapted from Fritz *et al.* [38], where fa is the population frequency of the g.4,039,268A allele, and μ is the phenotypic mean of the trait.

### Effects of heterozygosity for the Chr28 g.4,039,268A allele on performance traits

The Aubrac is one of the 9 breeds included in the French national genetic evaluation of beef cattle. Animals are genetically evaluated each year using the national polygenic BLUP evaluation for five traits measured in the commercial farms: birth weight, ease of calving, muscular development, skeletal development and weight at 210 days. Genetic breeding values and residuals were extracted from the French national database for genotyped animals and summed to obtain a phenotype adjusted for non-genetic effects. The effect of the g.4,039,268A allele was tested using the GWAS method for the five traits studied with the GCTA software version 1.26 [39], using the mlma option, and applied to the following mixed linear model:

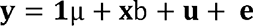

where y is the vector of corrected phenotypes; μ is the overall mean; l is a vector of ones; **b** is the additive effect of the derived allele; **x** is the genotype for the SNP; 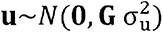 is the vector of random polygenic effect, where **G** is the genomic relationship matrix calculated using the 50K SNP genotypes (computed without Chr 28), and 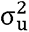 is the polygenic variance that is estimated based on the null model without the SNP effect; and 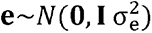 is the vector of random residual effects, where I the identity matrix and 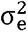 the residual variance. The number of animals analyzed ranged from 4,401 to 8,416 depending on the trait, of which 44% had a status based on direct genotyping of the variant and 56% based on a haplotype test considering the 35-marker haplotype (from positions 3,583,342 bp to 5,092,017 bp on the bovine reference genome assembly ARS-UCD1.2 [24]) identified by homozygosity mapping (see Results).

## Results

### Pedigree analysis suggests an autosomal recessive mode of inheritance

The cases consisted of eight males and thirteen females born to unaffected parents over an 18-year period in 21 purebred Aubrac herds spread throughout France. Analysis of the pedigrees of 17 cases with information available back to the 1960’s revealed several recent inbreeding loops supporting an autosomal recessive mode of inheritance, but did not allow us to identify a single ancestor shared by all their parents. Further analysis highlighted the bull “E.” (born in 1989) as the most influential spreader of this putative recessive defect in recent decades, with a ratio of 2.04 between its genetic contributions to the case group (3.20 %) and to 110,247 controls born between 2019 and 2021 (1.57%; Fig. 1 a). This AI bull was present in the genealogy of 12/17 cases (Fig. 1 b).

**Figure 1.**
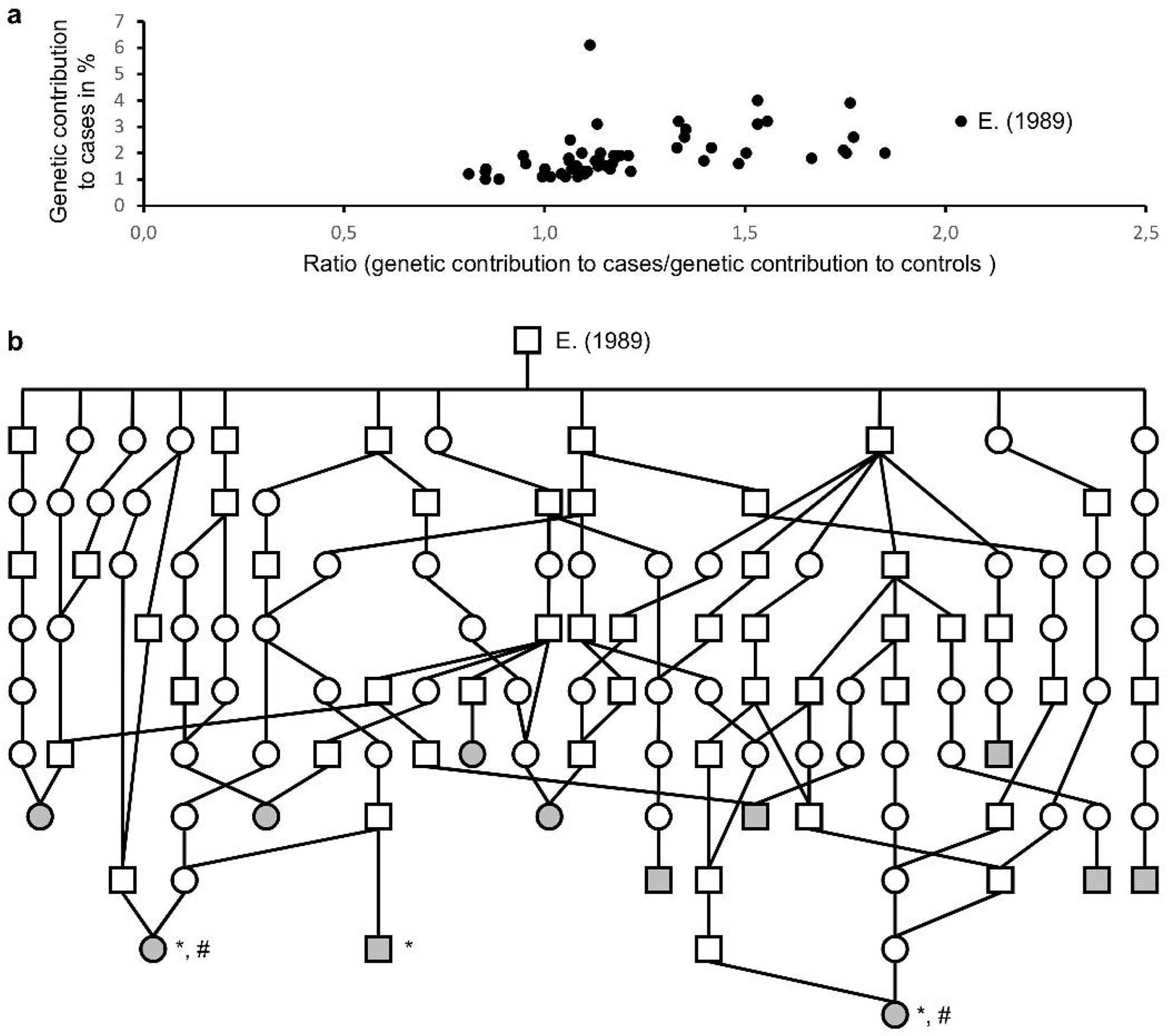
Pedigree analysis. (**a**) Graph showing the genetic contribution to the case group (n=17) and the ratio “contribution to the cases / contribution to 110,247 controls” for ancestors with a genetic contribution greater than or equal to 1% in each population. (**b**) Simplified pedigree of 12 cases descending from ancestor « E. ». Squares and circles represent males and females respectively. Gray filled symbols correspond to affected calves. *: Animals necropsied. #: Individuals selected for whole genome sequencing.

### Clinical findings are compatible with RCDP

The 21 affected calves were stillborn and exhibited extremely disproportionate dwarfism characterized by craniofacial dysmorphism, short limbs with hypermobile joints, a distended abdomen prone to eventration, and low birth weight despite normal gestation length (i.e., ∼20-30 kg versus ∼40 kg; Fig. 2). Due to the rapid collection of dead animals by rendering companies, only three affected calves (two females, one male) were available for extensive pathological examination. Radiographs, CT scans, and longitudinal skull sections allowed better characterization of the craniofacial dysmorphism. The latter consisted primarily of severe hypoplasia of the maxilla and secondary deformities of neighbouring bones and soft tissue structures, resulting in a cleft palate, curvature of the mandible, protrusion of the tongue, bossing of the frontal bone, and the presence of an anterior fontanelle (Fig. 2). Imaging and skeletal preparation also revealed platyspondyly of the thoracic and lumbar vertebrae, abnormally short ribs, and rhizomelic limb shortening (Additional file 2: Figure S1; Fig. 3). More specifically, the proximal long bones had shortened diaphyses and enlarged metaphyses with thickened cortex, whereas the diaphyses of the distal long bones (metatarsus, metacarpus, and phalanges) were normally developed. In addition, the tuberosity of the calcaneus, the femoral head and all epiphyses were absent or reduced to punctate calcifications (Fig. 3). Finally, the necropsy revealed hyperlaxity of all joints except the stifle and hock, which were affected by arthrogryposis, and no particular malformations of the internal organs.

**Figure 2.**
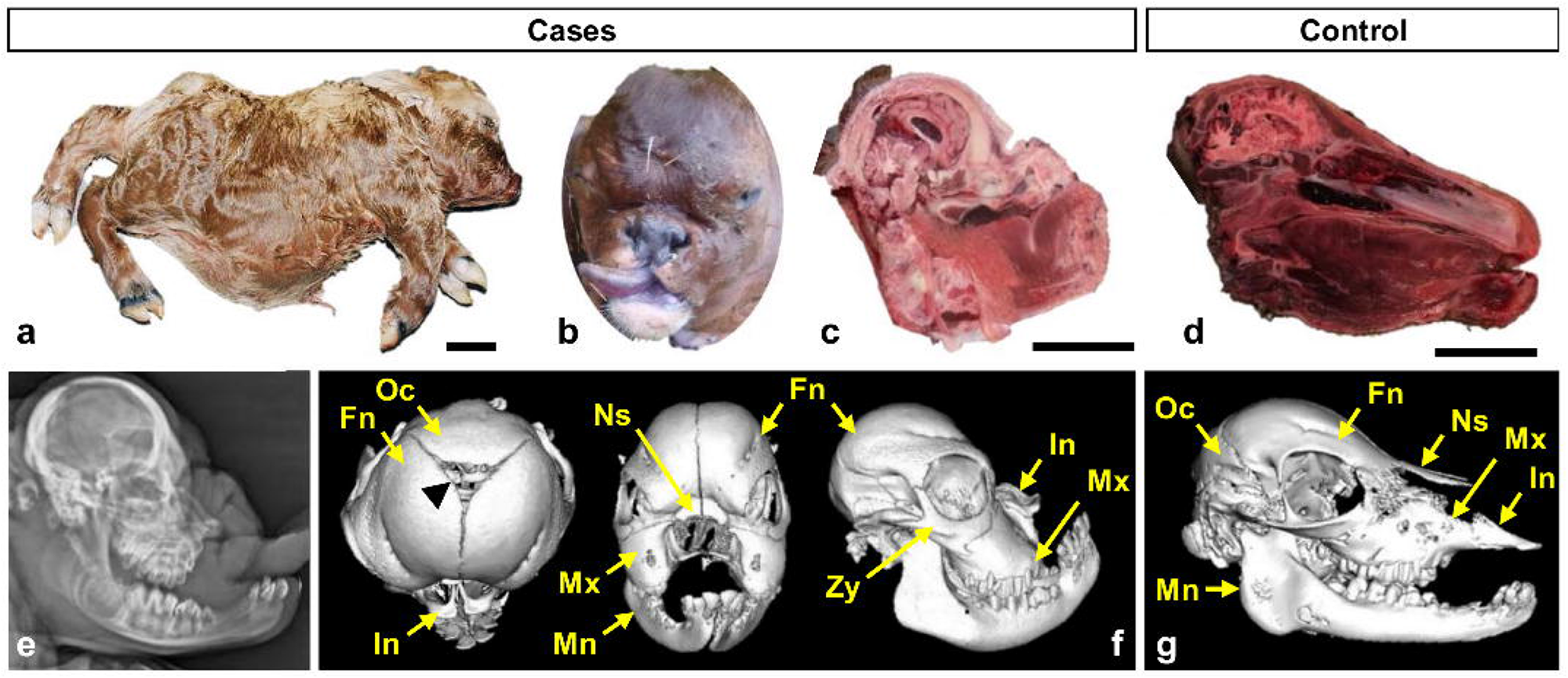
Macroscopic view of Aubrac « Bulldog » calves and characterization of their craniofacial dysmorphism. (**a-b**) General view and detail of the head of affected calves with facial features reminiscent of those of the French bulldog, hence the name given to this pathology by breeders. (**c-d**) Longitudinal section of the skull of a case and control calf, respectively. **e**) Radiograph of the head of an affected calf. (**f-g**) CT scan images of the head of a case and a control calf, respectively. The black arrowhead points to the anterior fontanelle between the occipital bone and the two frontal bones observed in affected calves. Fn: Frontal bone. In: Incisive bone. Mn: Mandible. Mx: Maxillary bone. Ns: Nasal bone. Oc: Occipital bone. Zy: Zygomatic bone. Scale bars = 10 cm.

**Figure 3.**
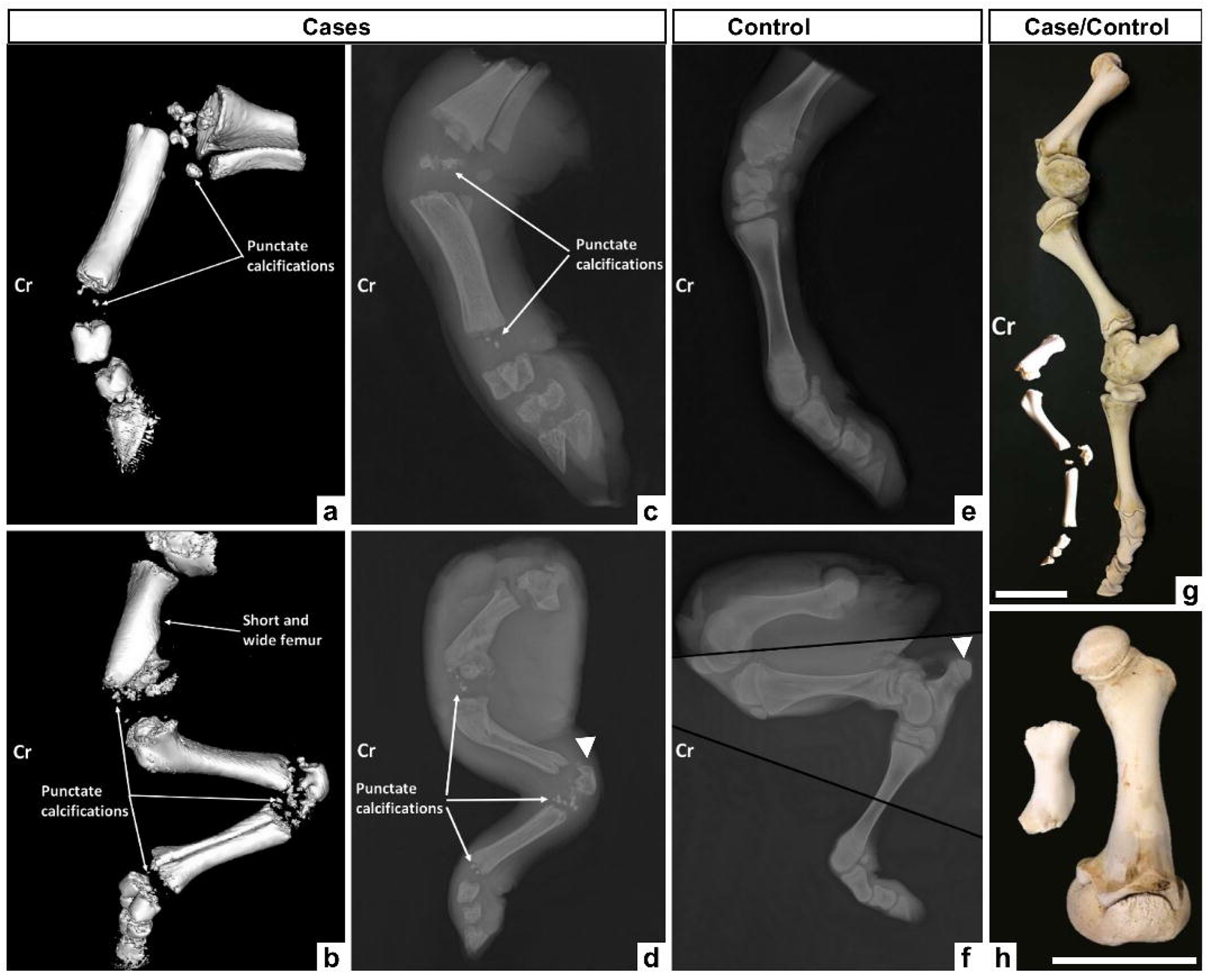
Imaging and skeletal preparation of the limbs of “Bulldog” and control calves. (**a-b**) CT scans of the anterior (**a**) and posterior (**b**) limbs of the case. (**c-d**) Radiographs of the limbs of the same RCDP affected individual. (**e-f**) Radiographs of the anterior (**e**) and posterior (**f**) limbs of a control individual. **g**) Skeletal preparation of the left hind limb of case and control calves. (**h**) Detail of the humerus shown in (**g**). Note the presence of multiple punctate calcifications where the epiphyses should be found and the absence of the femoral head (visible in (**h**)) and the tuberosity of the calcaneus (indicated by a white arrowhead in (**d**) and (**f**)). Cr: Cranial orientation. Scale bars = 10 cm.

Based on all of these elements, we arrived at the diagnosis of rhizomelic chondrodysplasia punctata (RCDP).

### Mapping and identification of a candidate causal variant in *GNPAT*

As a first step to gain insight into the molecular etiology of this bovine form of RCDP, we used a homozygosity mapping approach. By analyzing Illumina BovineSNP50 genotypes of 21 case and 1628 control animals for sliding windows of 20 markers, we mapped the RCDP locus at the beginning of chromosome 28 (Fig. 4 a). Under the peak position, we identified a 35-marker haplotype (from positions 3,583,342 bp to 5,092,017 bp on the bovine reference genome assembly ARS-UCD1.2) that was observed in the homozygous state in all the affected animals and in none of the controls. The most proximal markers outside of this segment defined the borders of a 1.6 Mb mapping interval (Chr28:3,555,723-5,143,700 bp) containing *GNPAT* and 11 additional genes (*Homo C1orf198, TTC13, ARV1, FAM89A, TRIM67, Homo C1orf131, EXOC8, SPRTN, EGLN1, TNSAX, DISC1*; Fig. 4 b).

**Figure 4.**
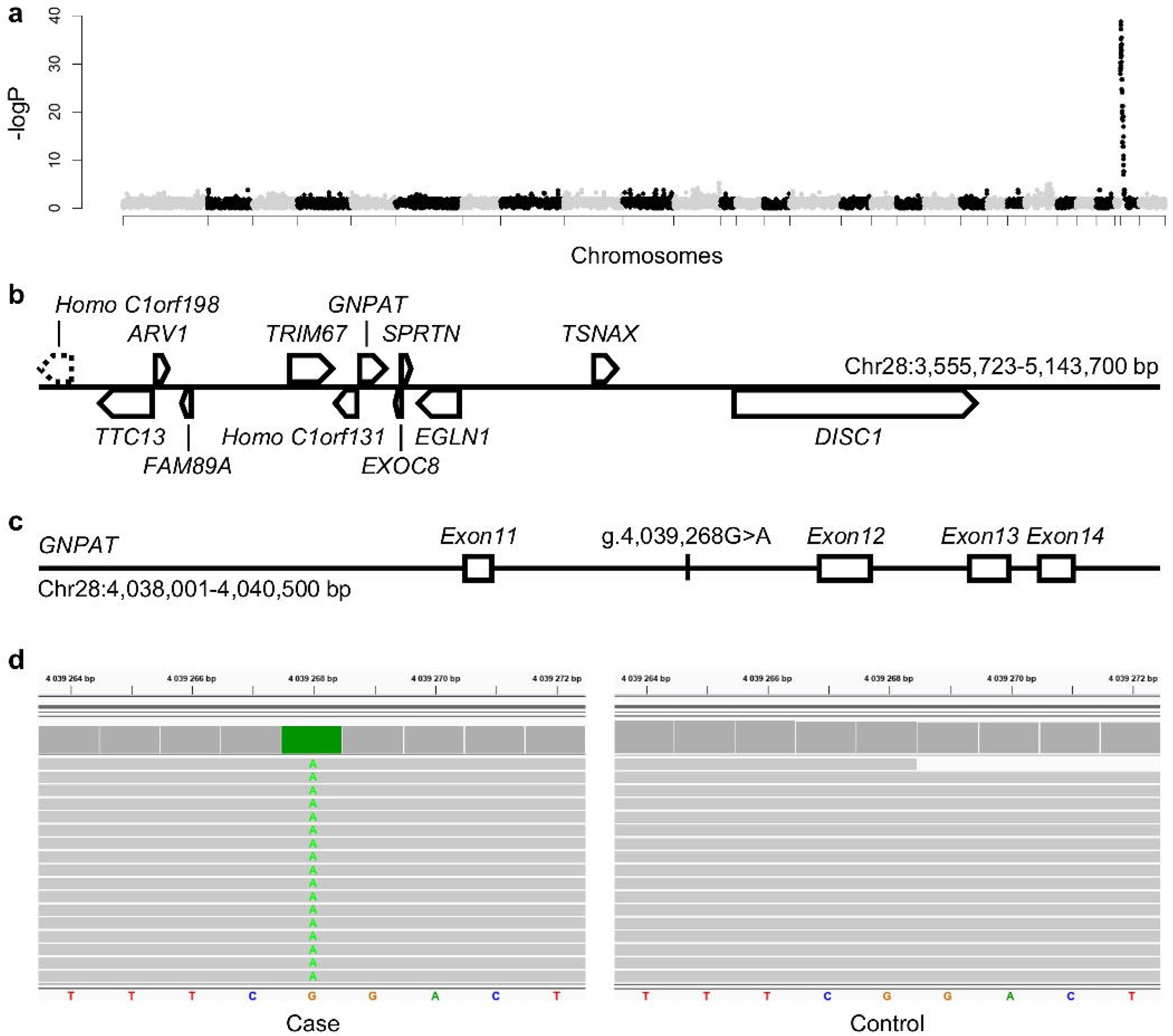
Mapping and identification of a candidate causative mutation in *GNPAT* intron 11. (**a**) Manhattan plot of homozygosity mapping results with phased and imputed Illumina BovineSNP50 array genotypes from 21 RDCP-affected and 1548 control individuals. The 0.05 significance threshold was set at −log P=6.20 after Bonferroni correction of Fisher’s exact test p-values. (**b**) Details of the genes located within the mapping interval. The dashed line indicates that the Homo C1orf198 gene encompasses the left border of the interval. (**c**) Details of the localization of the g.4,039,268G>A candidate variant in the 11^th^ intron of *GNPAT*. (**d**) Integrative Genomic Viewer screenshot showing homozygosity for this substitution in the whole genome sequence of an affected calf compared to a control.

Next, we sequenced the complete genomes of two RCDP-affected calves and compared them with 1,867 control genomes (Additional file 1: Table S1). The latter consisted of 39 non-carrier Aubrac individuals (based on haplotype information) as well as representatives of more than 70 breeds. Within the mapping interval, we identified a total of 3,115 sequence variants for which both cases were homozygous for the alternative allele (Additional file 3: Table S2). Subsequent filtering for variants completely absent in the controls reduced the list to a single candidate: a deep intronic substitution located 549 bp downstream and 323 bp upstream of *GNPAT* exons 11 and 12, respectively (Chr28 g.4,039,268G>A; Fig. 4 c, d).

### Validation of the *GNPAT* g.4,039,268G>A variant by large-scale genotyping

As a first verification, we genotyped the *GNPAT* g.4,039,268G>A variant in the 21 cases, all their available parents (n=26), and the AI bull “E.” using the Illumina EuroGMD custom SNP array. As expected, each case was homozygous for the derived allele while all unaffected parents and “E.” were heterozygous. For further validation, we extended the analysis to 1,195 unaffected Aubrac cattle and 375,535 controls from 17 breeds genotyped on the same array for genomic evaluation purposes. The g.4,039,268A allele was found to segregate only in Aubrac cattle at a frequency of 2.60% and we did not observe any homozygous carriers (Table 1).

**Table 1.**
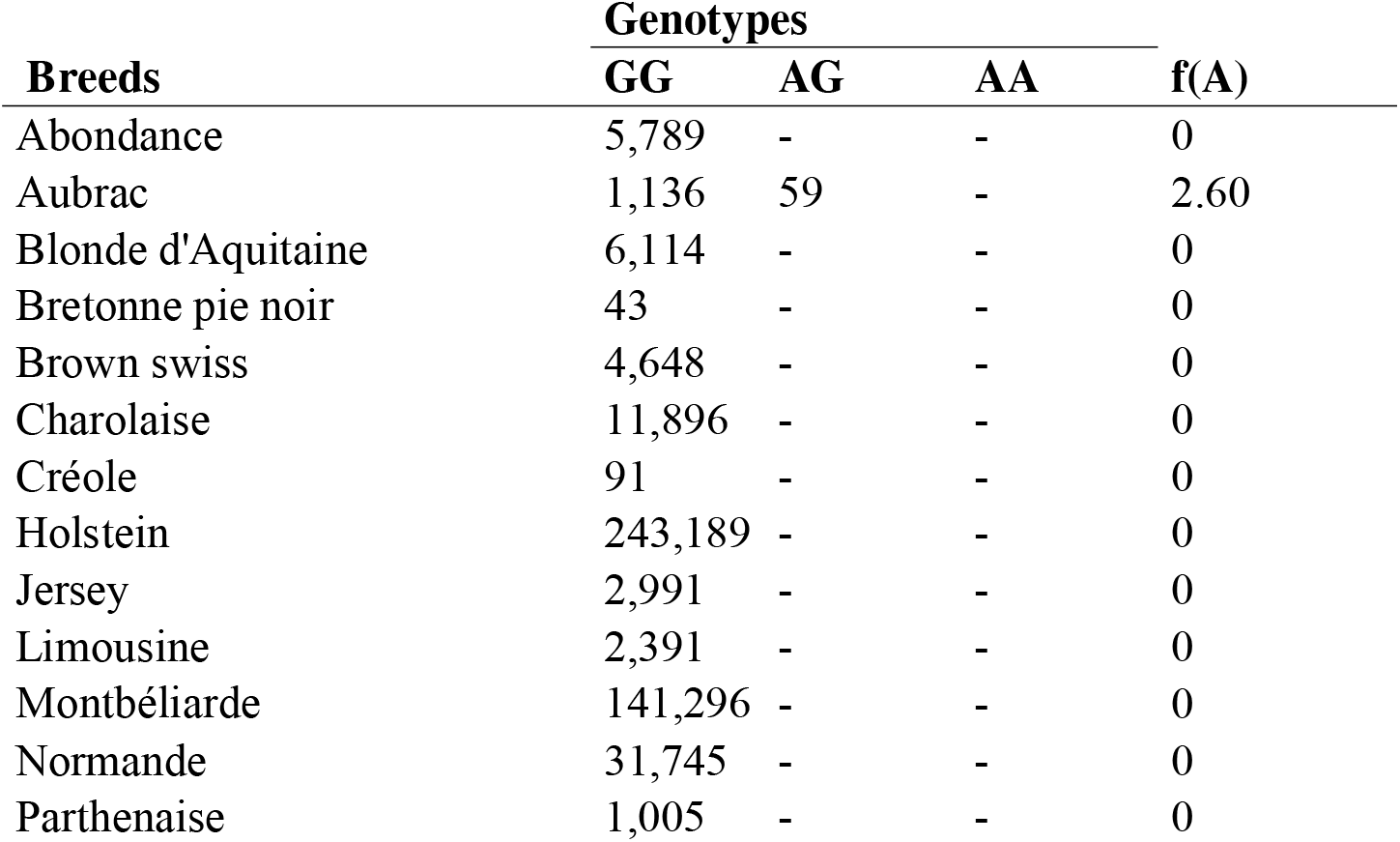
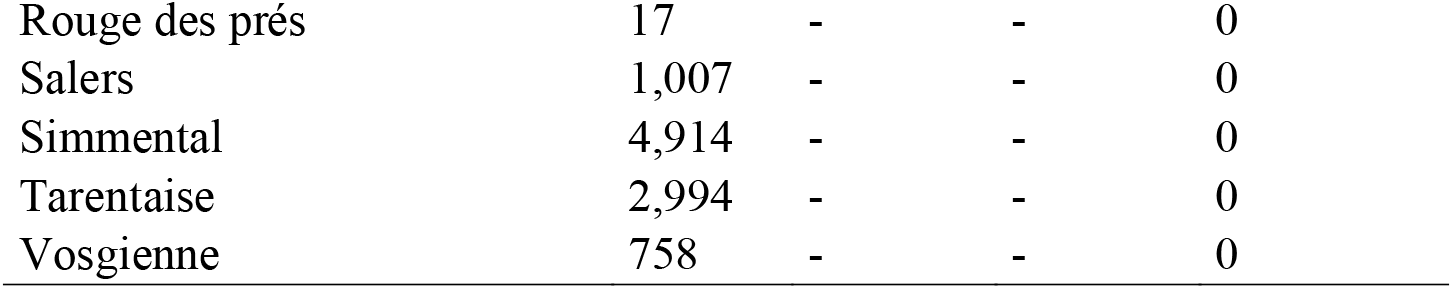
Results of large-scale genotyping of the *GNPAT* g.4,039,268G>A variant in 376,730 unaffected animals from 18 breeds. The number of animals is given for each genotype per breed; f(A): frequency of the g.4,039,268A allele.

### Chr28 g.4,039,268A allele activates cryptic splice sites in *GNPAT* intron 11

Following these preliminary verifications, we performed a series of analyses to investigate the effects of the g.4,039,268A variant on *GNPAT* function.

Because tissues from RCDP-affected calves were collected and frozen at −20°C only prior to the discovery of the candidate variant, they were not available for a posteriori RNA extraction. Therefore, we attempted to perform a Western blot analysis using an antibody directed against the N-terminal region of the GNPAT protein (ab75060, Abcam).

Unfortunatly, we were unable to detect a signal of the expected molecular weight in wild-type samples, either with proteins extracted in our laboratory from the muscle of a control animal or with a commercial extract (BT-102, GENTAUR; results not shown). Although this antibody has been used successfully against both human and mouse GNPAT [40,41], we concluded that it does not work with the bovine orthologous protein.

After this unsuccessful attempt, we performed a minigene analysis to investigate *in vitro* a possible effect of the Chr28 g.4,039,268A allele on altering *GNPAT* splicing. We constructed two expression plasmids containing exon 11, intron 11 and exon 12 of the *GNPAT* gene and either the ancestral or the derived allele of the deep intronic variant (pcDNA3.1-GNPAT_G and pcDNA3.1-GNPAT_A, respectively; Fig. 5 a). RT-PCR analysis of HEK293T cells transfected with both minigenes showed only one major specific transcript for each construction, but of different sizes (Fig. 5 a). Sanger sequencing of the amplicons revealed that transcript No. 1 from pcDNA3.1-GNPAT_G was fully spliced and corresponded to exon 11/exon 12 whereas transcript No. 2 from pcDNA3.1-GNPAT_A corresponded to exon 11 and exon 12 separated by a small portion of intron 11 consisting of an 86 bp cryptic exon (Chr28:4,039,260-4,039,345; Fig. 5 b, c). Consistent with this observation, sequence analysis using the ESEfinder 3.0 software revealed that allele A was predicted to increase the binding capacity of the SF2/ASF splicing factor at the 5’ end of the cryptic exon, which may explain the selective inclusion of the latter in transcript No. 2 (Fig. 5 c). As a final step, we performed RT-PCR analyses on total blood RNA extracted from three heterozygous (HT) and three wild-type (WT) Aubrac cattle to verify the effects of the deep intronic variant *in vivo*. cDNA amplification with primers targeting *GNPAT* exons 11 and 12, followed by agarose gel electrophoresis, yielded four distinct bands that were purified and sequenced by Sanger’s method (Fig. 5 d). Bands 1 and 4 were observed in both WT and HT animals and corresponded to amplicons with exon 11 and 12, and either fully spliced or unspliced intron 11, respectively. In contrast, bands 2 and 3 were observed exclusively in HT animals and resulted from abnormal splicing of intron 11. Between exons 11 and 12, PCR product 2 contained the same 86 bp cryptic exon observed in minigene analysis, whereas PCR product 3 also contained the portion of the intron 11 located between exon 11 and the cryptic exon, in addition to the latter.

**Figure 5.**
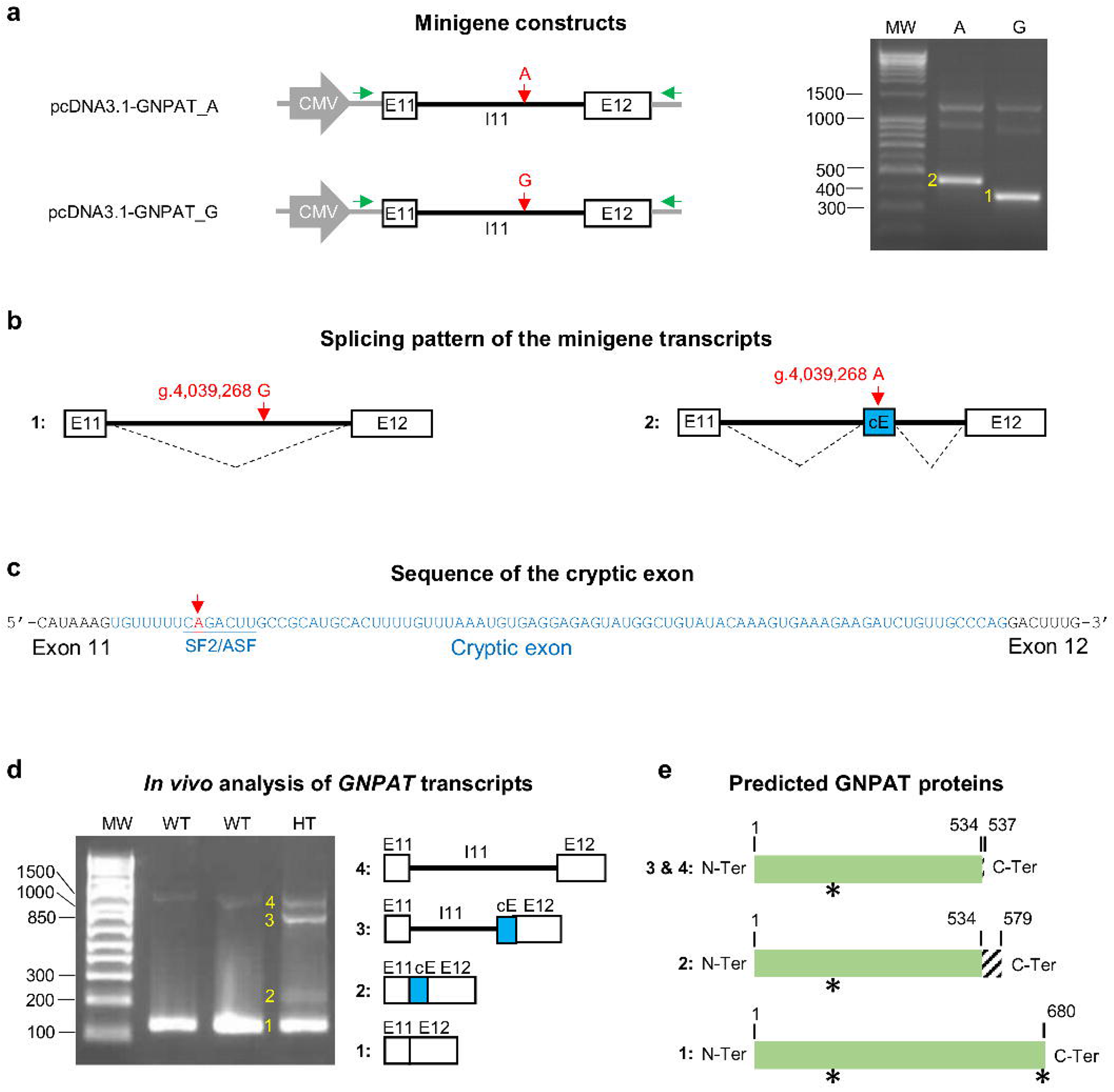
*In vitro, in silico* and *in vivo* analysis of the effect of the g.4,039,268G>A substitution on the *GNPAT* splicing. **(a)** Minigene analysis: Illustration of the pcDNA3.1-GNPAT minigenes carrying the derived or ancestral alleles in GNPAT intron 11 (upper and lower left panels, respectively) and results of the RT-PCR and gel electrophoresis after transfection of HEK293T cells with the pcDNA3.1-GNPAT_A (A) and pcDNA3.1-GNPAT_G (G) minigenes (right panel). cDNAs were amplified with primers targeting the pcDNA3.1 5’UTR and 3’UTR regions (green arrows). Two major PCR products were detected (labeled 1 and 2). MW: Molecular weight. **(b)** Splicing patterns associated with the two minigenes based on Sanger sequencing of the amplicons shown in (**a**). cE: Cryptic exon. (**c**) Sequence details of the cryptic exon: The g.4,039,268G>A substitution increases the score for a predicted SF2/ASF binding site located in its 5’ region according to the ESEfinder 3.0 software. (**d**) *In vivo* analysis of *GNPAT* transcripts. Left panel, representative subset of the results obtained after gel electrophoresis following RT-PCR on total blood RNA extracted from wild-type (WT) and heterozygous (HT) Aubrac cattle using primers targeting GNPAT exons 11 and 12. Four observed PCR products are numbered and their structures are shown (see text for details). MW: Molecular weight. (**e**) Consequences of the splicing patterns shown in (**d**) on the primary structure of the GNPAT protein. Normal amino acids (AAs) are shown in green, novel AAs are shaded. The acyltransferase motif (AAs 162 to 167) and the peroxysomal targeting signal 1 (PTS1, AAs 678 to 680; (18)) are marked with an asterisk.

Taken together, the results of our *in vitro*, *in silico* and *in vivo* analyses support that the Chr28 g.4039268A allele alters the *GNPAT* splicing by activating cryptic splice sites within intron 11. Incorporation of all or part of the latter intron into the *GNPAT* mRNA is predicted to cause frameshifts and to generate mutant proteins lacking the last 21% amino acids of the bovine GNPAT protein and, in particular, the C-terminal microbody targeting signal (Fig. 5 e).

### Mining the large dataset of records from the French national bovine database to study the effects of the Chr8 g.4,039,268A allele

As a final step to complete our study, we mined the large dataset of records from the French national bovine database to gain further insight into the phenotypic consequences of the Chr28 g.4,039,268A allele in the heterozygous or homozygous states.

First, we investigated the penetrance and expressivity of this allele by examining four juvenile mortality rates for different types of matings between genotyped sires and daughters of genotyped sires using a fixed effect model (Table 2). We observed a significant increase in mortality rates in at-risk (i.e., where the sire and maternal grandsire are both heterozygous; 1 x 1) versus control (wild-type sire and maternal grandsire; 0 x 0) matings for the 0-2 day period (+5.69 %; p-value <0.0001) and also for the 3-14 day period (+1.28%; p-value=0.02), indicating that a fraction of homozygous mutant calves were not stillborn but died a few days later. Combining these periods, the increase in mortality reached +6.89% within the first 2 weeks after birth (p-value <0.0001), which is only about half of the +12.30% increase in mortality expected in at-risk matings assuming complete penetrance (see Methods for calculation details). To assess whether this difference between the expected and observed increase in the mortality rate was due to incomplete penetrance or to underreporting of stillbirths, we examined the 21 cases reported to the ONAB and found that only 61.90% (13/21) of them had been ear-tagged and officially registered in the French national bovine database. Considering that these two proportions (observed/expected increase in mortality within the first 2 weeks= +6.89 %/+12.30%=56.02% and ear-tagged individuals/cases reported=61.90%) were not significantly different among 21 individuals using a Chi2 goodness of fit test (12:9 vs 13:8; p=1), we concluded that the penetrance of peri- and postnatal mortality is most likely complete in homozygous carriers of the Chr8 g.4,039,268A allele.

**Table 2.** Analysis of two juvenile mortality rates for different mating types for the Chr28 g.4,039,268G>A variant. “Mating” indicates the genotype for the Chr8 g.4,039,268G>A variant in allelic dosage (1=heterozygous carrier of the mutant allele; 0= non-carrier) of the sire and maternal grandsire of the group of individuals considered. For example, “1 x 0” refers to the progeny of a carrier bull with the daughter of a non-carrier bull. Nb: Number of observations. SE: Standard error. Difference (%): Difference between the studied mating type and the control group (i.e., mating type 0 x 0). P-value: Student’s t-test. Note the significant differences between the 1 x 1 and 0 x 0 genotype groups for several periods. Note also that we found a small but significant +0.47 increase in mortality for the 0-2 day period in matings between carrier sires and non-carrier sires (1 x 0 vs. 0 x 0) due to the fact that the mutant allele segregates at a frequency of 2.60% in the maternal granddam population.

| Trait | Mating | Nb | Raw mean (%) | Difference (%) | SE (%) | P-value |
| --- | --- | --- | --- | --- | --- | --- |
| 56-365 days mortality rate | 1 x 1 | 268 | 1.49 | 0.47 | 0.62 | 0.44 |
|  | 1 x 0 | 4078 | 1.08 | 0.06 | 0.16 | 0.72 |
|  | 0 x 1 | 4973 | 0.95 | -0.08 | 0.15 | 0.60 |
|  | 0 x 0 | 91703 | 1.02 | - | - | - |
| 15-55 days mortality rate | 1 x 1 | 271 | 1.11 | 0.37 | 0.53 | 0.48 |
|  | 1 x 0 | 4,111 | 0.80 | 0.07 | 0.14 | 0.63 |
|  | 0 x 1 | 5,022 | 0.94 | 0.20 | 0.13 | 0.11 |
|  | 0 x 0 | 92,397 | 0.74 | - | - | - |
| 3-14 days mortality rate | 1 x 1 | 277 | 2.17 | 1.28 | 0.57 | 0.02 |
|  | 1 x 0 | 4,144 | 0.80 | -0.09 | 0.15 | 0.54 |
|  | 0 x 1 | 5,077 | 1.08 | 0.20 | 0.14 | 0.15 |
|  | 0 x 0 | 93,227 | 0.89 | - | - | - |
| 0-2 days mortality rate | 1 x 1 | 300 | 7.67 | 5.69 | 0.81 | 1.59*10<sup>-11</sup> |
|  | 1 x 0 | 4,248 | 2.45 | 0.47 | 0.22 | 0.03 |
|  | 0 x 1 | 5,178 | 1.95 | -0.03 | 0.20 | 0.89 |
|  | 0 x 0 | 95,109 | 1.98 | - | - | - |
| **0-14 days mortality rate** | **1 x 1** | **300** | **9.67** | **6.82** | **0.97** | **1.32\*10<sup>-11</sup>** |
|  | 1 x 0 | 4,248 | 3.23 | 0.38 | 0.26 | 0.15 |
|  | 0 x 1 | 5,178 | 3.01 | 0.16 | 0.24 | 0.49 |
|  | 0 x 0 | 95,109 | 2.85 | - | - | - |

Finally, we studied the effects of the Chr28 g.4,039,268A allele on five performance traits genetically evaluated each year in the framework of the national polygenic BLUP evaluation. We found a significant result for only one trait, namely a reduction of one point of muscular development (MDev) at the age of 210 days in heterozygous carriers versus wild-type inviduals (p=0.04; Table 3). Note that 1.00 point represents 21% of the genetic standard deviation for this trait (GSD=4.77 points).

**Table 3:** Analysis of five performance traits in animals genotyped for Chr28 g.4,039,268G>A variant. MDev: Muscular development. SDev: Skeletal development. B effect: effect size. SD: Standard deviation. P-value: Student’s t-test.

| Trait | Headcount per genotype |  | B effect (SD) | P-value |
| --- | --- | --- | --- | --- |
|  | Wild-type | Heterozygous |  |  |
| Birth weight (in kg) | 7,928 | 488 | -0.08 (0.13) | 0.54 |
| Ease of calving (score from 1 to 5) | 7,926 | 488 | 0.00 (0.01) | 0.91 |
| MDev at 210 days (score on 100) | 4,404 | 271 | -1.00 (0.49) | **0.04** |
| SDev at 210 days (score on 100) | 4,404 | 271 | 0.24 (0.48) | 0.61 |
| Weight at 210 days (in kg) | 4,146 | 255 | -0.71 (1.14) | 0.53 |

## Discussion

In this article, we set up a powerful approach that led to the identification and characterization of a novel *GNPAT* deep intronic splicing mutation responsible for recessive RCDP in Aubrac cattle. The success of our strategy largely relied on the particular structure of the bovine populations and on the availability of massive amounts of pedigree, genomic and phenotypic information recorded for selection purposes [5].

Cattle breeds are genetically small populations created 150 years ago from a limited number of founders, whose genetic variability has been further reduced over the last 50 years by the overuse of influential sires through AI [42,43]. Typically, cattle breeds have effective population sizes (Ne) ranging from 12 to 150, and a minimum number of ancestors contributing to 50% of the breed’s gene pool ranging from 5 to 71, as reported in a recent study of 26 cattle breeds reared in France (https://idele.fr/detail-dossier/varume-resultats-2023; accessed 2024/03/28). This low genetic variability within breeds contrasts with the very high genetic variability observed at the species level, supported by the discovery in 2019 of 84 million SNPs and 2.5 million small insertion-deletions (equivalent to one variant every 31 bp) in a collection of 2,703 individuals representing a significant proportion of global cattle population diversity as part of the 1000 Bull Genomes Project [44].

As a result, it is possible to capture most of a breed’s gene pool by sequencing only its major ancestors (for whom biological material is often available decades after their death in the form of frozen semen straws), and to capture ancient genetic variability by repeating this sequencing effort in numerous independent breeds. This situation explains why, after homozygosity mapping of the RCDP locus in a 1.6-Mb interval on Chr28, we were able to reduce the number of candidate variants from 3,115 to a single one: a deep intronic substitution located in the 11^th^ intron of the *GNPAT* gene (Chr28 g.4039268G>A).

Although this type of variant usually does not affect gene expression, in rare situations it can cause severe splicing defects, often resulting in monogenic defects due to partial or complete loss of gene function (e.g. [45,46]). To our knowledge, the only example reported so far in cattle is a SNP in intron 2 of the gene encoding myostatin (MSTN), which causes muscle hypertrophy in the Blonde d’Aquitaine breed [47]. As several loss-of-function mutations in the coding part of *MSTN* had been previously described in double-muscled cattle [48,49], the authors used a candidate gene approach: they first sequenced the cDNA of this gene and then its unique exon, after observing an abnormal transcript.

Here, since the Chr28 g.4039268G>A substitution was the only remaining candidate variant after a thorough mapping, whole genome sequencing and filtering procedure, we logically assumed that it altered the correct splicing of *GNPAT* and performed complementary *in vitro*, *in vivo, and in silico* analyses to validate this hypothesis.

As RNA is rapidly degraded at room temperature within a few hours of death, we could not study the expression of *GNPAT* in homozygous mutants. We therefore chose to analyze the splicing of two minigenes (containing intron 11 with either the ancestral or the derived allele of the Chr28 g.4039268G>A substitution and the two flanking exons) in HEK293T transfected cells and, subsequently, of *GNPAT* in blood RNA samples from three heterozygous carriers and three wild-type Aubrac cattle (Fig. 5).

Both confirmed that the derived allele was associated with abnormal splicing patterns, likely mediated by an increase in the binding capacity of an SF2/ASF splicing factor encompassing the deep intronic substitution, as predicted by the ESEfinder 3.0 software.

The *in vitro* analysis yielded a transcript containing an 86 bp cryptic exon, which was also observed *in vivo*, together with two additional aberrant transcripts corresponding to incomplete splicing of the newly created introns surrounding this cryptic exon.

However, while the level of expression of the two minigene constructs were similar *in vitro*, the aberrant transcripts appeared to be less abundant than the correctly spliced ones in the blood of heterozygous animals. We suggest that this difference is due to degradation of the mispliced transcripts containing premature termination codons (PTC) by non-sense-mediated mRNA decay (NMD) *in vivo* but not *in vitro*. In fact, NMD cannot occur in the context of the minigene because the PTC is located in the last exon, which is not the case in the context of the full gene sequence *in vivo*.

In any event, translation of transcripts containing all or part of intron 11 would result in the production of proteins lacking the last 21% amino acids of the bovine GNPAT protein (Figure 5e), and in particular the peroxysomal targeting signal 1, which is essential for sorting the majority of peroxysomal proteins to this organelle [50,51].

Therefore, whether due to NMD or to frameshifts and targeting errors, our results suggest that the splicing defects caused by the Chr28 g.4039268G>A deep intronic mutation will ultimately lead to a major reduction in the amount of functional GNPAT protein in the peroxysomes of homozygous mutants.

The glycerone-phosphate O-acyltransferase (GNPAT), also known as dihydroxyacetone phosphate acyltransferase (DAP-AT, DAPAT and DHAPAT), is an enzyme located exclusively in the peroxisomal membrane that mediates the first step in the synthesis of ether phospholipids including plasmalogen [52].

As for other proteins involved in peroxisomal protein import (PEX5 and PEX7) or ether phospholipid synthesis (AGPS and FAR1), mutations in the gene encoding GNPAT have been reported to cause RCDP in humans [15–20].

RCDP is a severe developmental disorder caused by defects in plasmalogen synthesis. Patients with RCDP present with skeletal dysplasia including rhizomelic shortening of the limbs, characteristic punctate epiphyseal calcifications, and a typical dysmorphic facial appearance with a broad nasal bridge, epicanthus, high-arched palate, micrognathia, and dysplastic external ears. Clinical features also include congenital cataracts, contractures, seizures, severe growth and psychomotor retardation, and markedly shortened life span [16,53,54]. The degree of plasmalogen deficiency, which depends on the gene and type of mutation involved, determines the severity of the syndrome [55]. For example, erythrocyte plasmalogen levels are almost undetectable in classical severe RCDP, whereas they reached up to 43% of average controls in a study focused on 16 patients with mild RCDP [56]. To our knowledge, no deep intronic mutation in *GNPAT* has been reported in humans. However, based on a literature review, we identified three mutations that, similar to the bovine mutation reported here, are predicted to generate NMD-targeted mRNAs and proteins truncated in the C-terminal region (*GNPAT* c.1428delC, c.1483delG, and c.1575delC causing frameshifts starting at amino acid positions 477, 495, and 525, respectively; [57;58;17]. Plasmalogen levels measured in patients homozygous for any of these three deleterious variants were undetectable or close to zero, suggesting that they do not produce functional GNPAT proteins. Furthermore, neither GNPAT activity nor GNPAT protein was detected in cultured fibroblasts from patients with the *GNPAT*^c.1428delC/ c.1428delC^ and *GNPAT*^c.1483delG/ c.1483delG^ genotypes [57,58]. Here, because of some limitations due to the stillbirth and freezing of the necropsied specimens, we were not able to examine the neuromuscular manifestations or measure plasmalogen levels, which are traditionally evaluated in red blood cells by gas chromatography/mass spectrometry [59]. However, based on imaging and skeletal examination of three calves homozygous for the GNPAT deep intronic variant, we observed rhizomelic shortening of the limbs and punctate epiphyseal calcification, which are the hallmarks of the classic severe form of RCDP along with multiple craniofacial malformations. Thanks to the combination of large-scale genotyping and the mining of pedigree and performance records available in the French National Cattle Database, we also documented increased levels of juvenile mortality in the offspring of at-risk versus control mating (with some cases dying at birth and others within the following two weeks), consistent with full penetrance of the mutation in the homozygous state. Taken together, these observations further support our expectation that homozygosity for the bovine Chr28 g.4039268A allele results in severe GNPAT and plasmalogen insufficiency.

In addition, we would like to point out that, unlike the affected calves reported in this article, the clinical features of GNPAT-deficient mice are not entirely consistent with those commonly reported for human RCDP. Mice homozygous for a a targeted invalidation of the *Gnpat* gene exhibited a complete lack of plasmalogens, male infertility, defects in eye and central nervous system development, abnormal behavior, and mild skeletal abnormalities consisting of disproportionate dwarfism with shortening of the proximal limbs [60]. *Gnpat* KO mice were viable and while some of them died prematurely (∼40% within the first four to six weeks), others, especially females, were long-lived. This difference between humans and mice in the severity of clinical features associated with inactivation of a gene associated with RCDP was also observed for Pex7 (reviewed in [61]). This finding suggests that calves homozygous for the Chr28 g.4,039,268A allele could be used as a reliable large animal model for RCDP, as an alternative to mouse models, especially to study how impairment of plasmalogen biosynthesis may affect the process of endochondral ossification in this pathology.

The efficient identification of heterozygous carriers by genotyping of the GNPAT deep intronic mutation as part of the genomic evaluation, and the mastery of reproductive biotechnologies in livestock breeding make it possible to envisage the production of case and control individuals in experimental farms for further functional analyses and translational research between cattle and humans. Indeed, techniques such as oestrus synchronization, polyovulation, embryo collection, preimplantation diagnosis, embryo freezing and embryo transfer have been successfully used over the past two decades to study developmental processes such as horn ontogenesis in bovine fetuses (e.g., [62–64]).

Finally, mining performance records for thousands of genotyped cattle we report a significant reduction of muscular development at the age of 210 days in heterozygous carriers versus wild-type inviduals (p=0.04; n=271 and 4,404 individuals respectively). The magnitude of this reduction, which represents 21% of the genetic standard deviation for this trait, is similar to that of a QTL with a relatively high effect. This result is consistent with the observations of Dorninger et al. [65], that Gnpat KO mice have altered development and function of the neuromuscular junction, causing reduced muscle strength, and advocates for the fine phenotypic characterization of muscle development and function in humans heterozygous for GNPAT deleterious mutations.

## Conclusions

In conclusion, this study highlights the usefulness of large data sets available in cattle for (i) detecting causative mutations beyond the coding regions and (ii) characterizing their phenotypic effects, as exemplified by the report of the first large animal model of RCDP in humans caused by a deep intronic splicing mutation of *GNPAT*.

## Declarations

### Ethics approval and consent to participate

Experiments reported in this work comply with the ethical guidelines of the French National Research Institute for Agriculture, Food and Environment (INRAE). No permit for experimentation was required by law (European directive 2010/63/UE) since the affected animals were not purposedly generated for this study and since all invasive exams and sampling were performed post-mortem on animals that died of natural death. Blood was collected during routine sampling (for annual prophylaxis, paternity testing, or genomic selection purpose) by trained veterinarians and following standard procedures and relevant national guidelines. All the samples and data analyzed in the present study were obtained with the permission of the breeders and of the “OS Race Aubrac” breed organization.

### Consent for publication

Not applicable

### Availability of data and materials

The WGS data of the RCDP-affected calves are available at the European Nucleotide Archive (www.ebi.ac.uk/ena) under the study accession no. PRJEB76441.

## Authors’ contributions

A. Boulling and AC conceived and coordinated the project. JC, VP, AC, and JM performed necropsy and pathological examination. A. Barbat and CG analyzed pedigree information. CG supervised sample collection and preparation for SNP array genotyping and whole genome sequencing, and performed PCR and Sanger sequencing. AC analyzed SNP array genotypes. MB and AC analyzed whole-genome sequences. A. Boulling and LBB performed *in vitro*, *in silico* and *in vivo* analyses. AC contributed to *in silico* analyses. A. Barbat analyzed juvenile mortality. ST analyzed performance traits. HL, SF, CL, AD, RG and DB contributed reagents/materials/analysis tools. A. Boulling, AC and JC drafted the manuscript. All authors read and approved the final manuscript.

## Acknowledgments

We are grateful to Jacques Renou, director of the OS Aubrac breed organization and its staff, as well as the numerous veterinarians and Aubrac breeders involved in this project for providing samples, pedigree and phenotype information. We would also like to thank the INRAE GeT-PlaGE platform (http://get.genotoul.fr) for sequencing the genomes and our colleagues Nicolas Gaiani, Chris Hozé and Maya Lambert for their occasional assistance.

## Additional files

### Additional file 1: Table S1

Format: xlsx

Title: Details of the whole genome sequences used as controls in this study.

Description: See the following URLs for more information on the biosample and bioproject IDs: https://www.ncbi.nlm.nih.gov/biosample/ and https://www.ncbi.nlm.nih.gov/bioproject/. Nb_Ind_Breed: Number of individuals per breed.

### Additional file 2: Figure S1

Format: pdf

Title: Radiographs of an affected calf.

Description: Cr: cranial orientation. Scale bar = 10 cm.

### Additional file 3: Table S2

Format: xlsx

Title: List of homozygous positional candidate variants found in the genomes of two RDCP-affected calves

Description: Chr: Chromosome. “Present_in_controls” indicates whether the variant was observed in at least one of the 1,867 genomes used as controls (see Additional file 1: Table S1).

