## Supplementary figures and images for "A bovine model of rhizomelic chondrodysplasia punctata caused by a deep intronic splicing mutation in the *GNPAT* gene"

### Figure S1

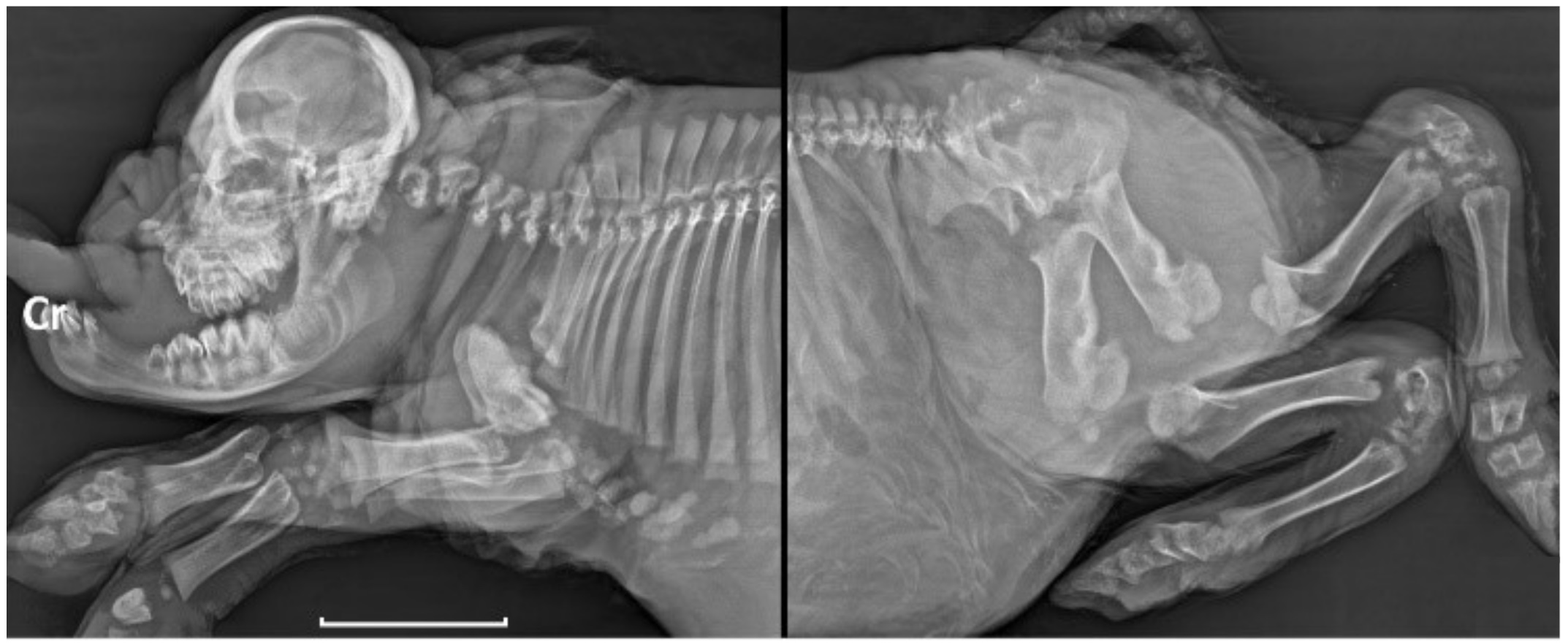
